# A semi-automatic dispenser for solid and liquid food in aquatic facilities

**DOI:** 10.1101/535062

**Authors:** Raphaël Candelier, Alex Bois, Stéphane Tronche, Jéremy Mahieu, Abdelkrim Mannioui

**Affiliations:** Sorbonne Université, Centre National de la Recherche Scientifique, Laboratoire Jean Perrin, LJP, F-75005 Paris, France; Sorbonne Université, Institut de Biologie Paris-Seine (IBPS), Aquatic Facility, 75005, Paris, France

## Abstract

We present a novel, low-footprint and low-cost semi-automatic system for delivering solid and liquid food to zebrafish, and more generally to aquatic animals raised in racks of tanks. It is composed of a portable main module equipped with a contactless reader that adjusts the quantity to deliver for each tank, and either a solid food module or a liquid food module. Solid food comprises virtually any kind of dry powder or grains below two millimeters in diameter, and, for liquid-mediated food, brine shrimps (*Artemia salina*) and rotifers (*Rotifera*) have been successfully tested. Real-world testing, feedback and validation have been performed in a zebrafish facility for several months. In comparison with manual feeding this system mitigates the appearance of musculoskeletal disorders among regularly-feeding staff, and let operators observe the animals’ behavior instead of being focused on quantities to deliver. We also tested the accuracy of both humans and our dispenser and found that the semi-automatic system is much more reliable, with respectively 7-fold and 84-fold drops in standard deviation for solid and liquid food.

## Introduction

Since the pioneering work of Streisinger *et al.* published in 1981 on cloning homozygous diploid zebrafish^1^, *Danio Rerio* has rapidly grown as a model system in many different fields including embryo development, tissue regeneration and neuroscience. In 2017, it has been estimated that more than 5 million zebrafish were used in more than 3.250 institutes spread across 100 countries^2,3^. It is more common in laboratories than other well-established aquatic vertebrates like xenopus^4^ and medaka^5^ and, for instance, it is the second most common animal species used for research in Great Britain^6^. As research on zebrafish has gained an impressive momentum in such a short time lapse, a large number of dedicated fish rooms have appeared. Other species like *Danionella Translucida* (now amenable to brain-wide functional imaging in adults with cellular resolution^7^) and killifish (whose short lifespan, fecundity and diapause of dried eggs make an ideal model for biogerontology^8^) also have a high potential for a rapid spread among research institutes in the future. Altogether, there has been and there will be a growing need for improving husbandry procedures in aquatic facilities, with very different scales ranging from a few hundred to hundreds of thousands animals.

Feeding is a fundamental task in any fish room, and though some research focus on improving the nutritional quality of the food^9,10^, only a few technical advances have been proposed on how food is actually delivered. It is traditionally performed manually with wash bottles for live food in liquid medium (*e.g. Artemia* nauplii, rotifers) and with various systems ranging from spoon-like tools to seed sowers for powders and granulates. Manual feeding raises serious issues though, with a clear lack of control over the delivered quantities and the appearance of musculoskeletal disorders among technicians. More specifically, wrist, elbow and shoulder tendinopathy are common among fish room staff, mainly because of the repetitive application of pressure on wash bottles. It may cause recurrent work stoppages, and in the most severe cases require steroid injections and surgery.

The only serious alternative to manual feeding is a fully-automated commercial solution, but it is extremely expensive, has a large footprint (which makes it difficult or impossible to install in small spaces, stand-alone racks or some fish rooms located in buildings that were not initially built for this purpose), processes very slowly (thus does not guarantee that microorganisms are still alive when delivered to the animals), delivers discrete quantities of food (which has limited accuracy), prevents access to the tanks during operation, and still requires food-filling and regular maintenance. In addition, breakdown – which is an inherent risk to every automated system – can have catastrophic consequences for both animals and the research associated.

Here we present an intermediate solution between manual and fully-automated systems, keeping the assets of both approaches while eliminating most of their drawbacks. Our Semi-Automatic Food Dispenser (SeAFooD) is battery-powered and portable with a low footprint, delivers dry solid or liquid-mediated food in a modular manner, displaces all the weight of liquid in a self-supporting reservoir on caster wheels, requires no specific gesture for triggering and remains under the operation of a human agent at all times. We quantified that microorganisms have a high or perfect survival rate while going through the dispenser and that survivors are as motile as control. The dispenser can deliver either fixed quantities, operator-controlled quantities or obtain information on the number of individuals in each tank *via* near-field communication (NFC) and automatically deliver the exact amount of food. The latter mode (*i*) has an accuracy down to the single-animal scale or below, (*ii*) diminishes waste and improves water quality, (*iii*) allows for custom diets and (*iv*) let the operator focus on animal behavior during the feeding process. The whole system is low-cost and has been built with standard tools anyone can find in a FabLab (*e.g.* 3D printer, laser-cutter, soldering iron). Finally, it has been tested in a medium-scale zebrafish platform for several months and had a very positive impact on the staff health since all staff members observed a decrease in forearm pains during this period.

## Materials and methods

### General description

The dispenser is composed of three modules (Figure 1, A): a *main module*, a *solid food module* for dry powders and grains and a *liquid food module* for live microorganisms in water. The main module has an ergonomic handle, a trigger, fixation rails to attach the other modules and a microcontroller to interface a LCD screen, a rotary encoder (to select the mode of operation and navigate in the settings menu), a NFC read/write card, and a high-power LED (Figure 2, A). The trigger has been designed to fit on Tecniplast tanks, but it can be easily commuted to fit other types of tanks. The main module always acts as the master while other modules are slaves.

**Figure 1.**
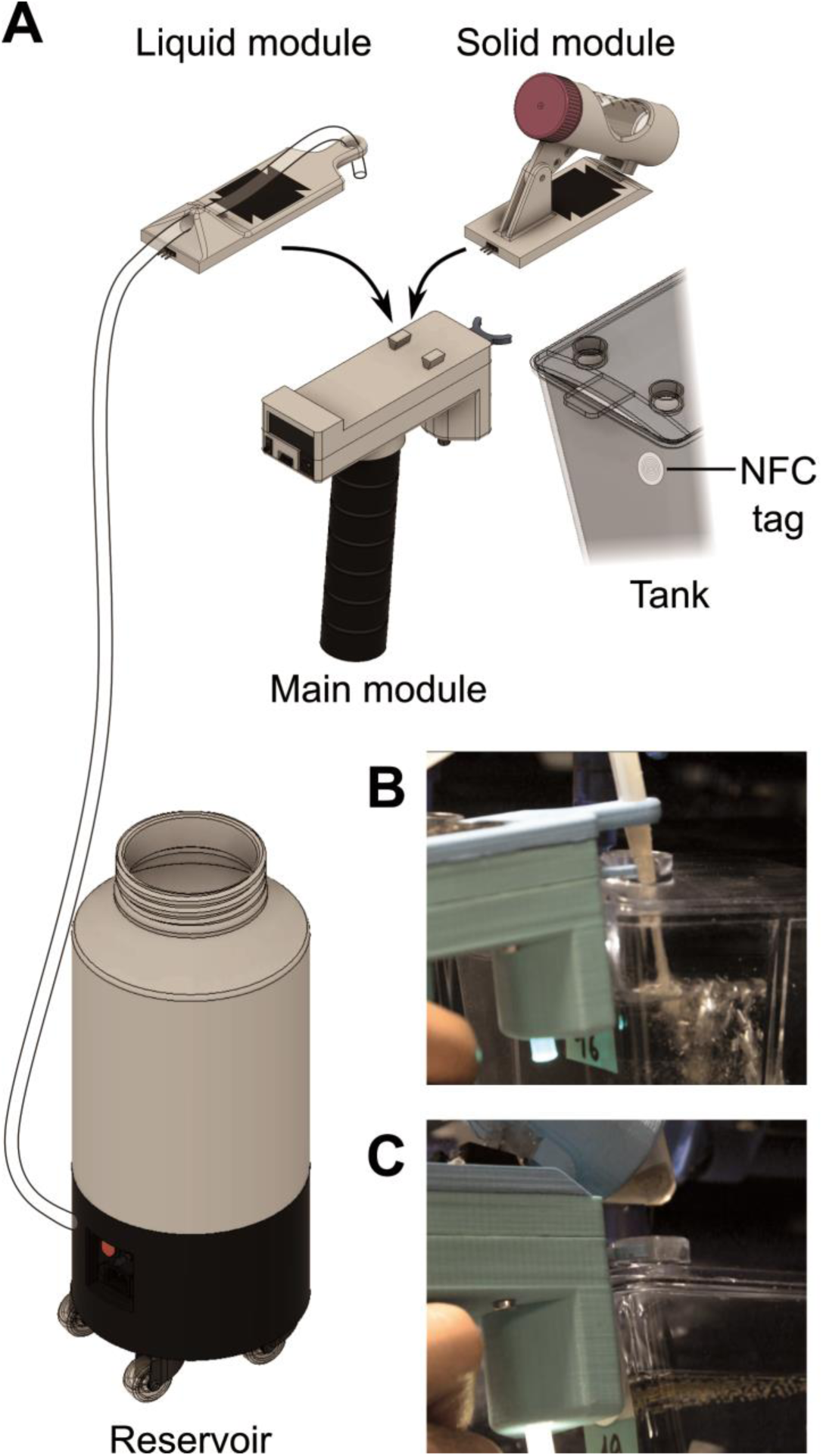
The semi-automatic food dispenser. (**A**) Scheme of the system. The main module can host either the liquid food or solid food module to form a functional assembly. The solid food module directly hosts 50mL tubes of powder while the liquid food module has an external 8L reservoir mounted on caster wheels. (**B**) Picture of the liquid food assembly during delivering. (**C**) Picture of the solid food assembly during delivering.

**Figure 2.**
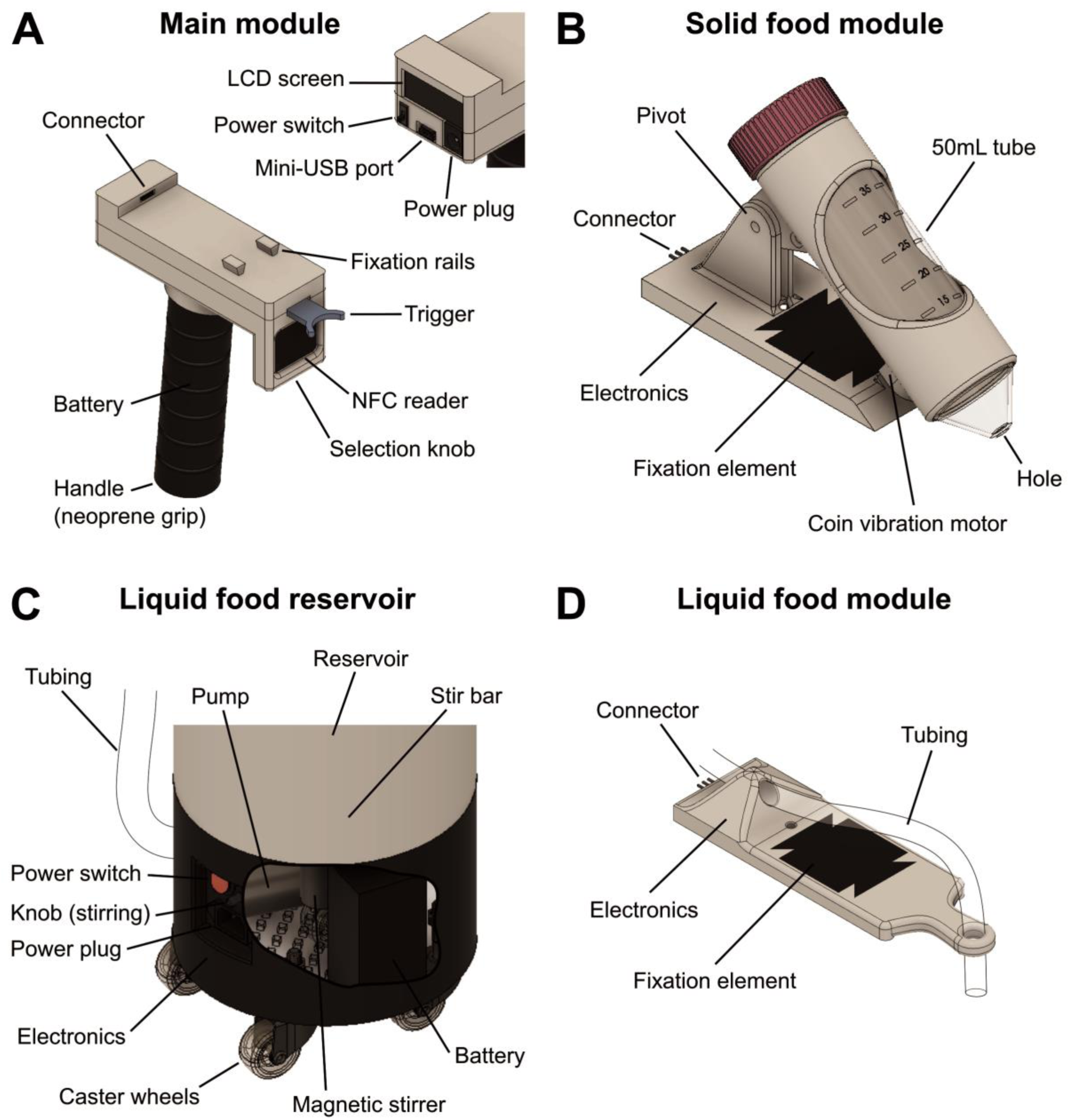
System parts and localization of relevant elements. (**A**) Main module. Inset: view of the back of the main module (**B**) Solid food module. (**C**) Zoom and partial inside view of the skirt of the liquid food reservoir. (**D**) Liquid food module.

The solid food module has a removable reservoir (standard 50mL tube, Falcon ref. 352070) drilled at the tip and mounted on a sheath with a vibration coin motor (Figure 2, B). During normal handling only minute amounts of powder smear from the reservoir, but under vibration a regular flow of grains instantaneously establishes (Supplementary Movie 1). This phenomenon has been previously described^11^ and, in essence, relies on the fluidization of the granular bed by a constant injection of energy to overcome friction among grains. Vibrations are similar to those of video games paddles, and are neither unpleasant nor dangerous for the operator.

The liquid food reservoir (8L) is attached onto a custom skirt with caster wheels (Figure 2, C). The skirt comprises a battery, a pump and a magnetic stirrer. The latter is essential for ensuring homogeneity in the solution and *Artemia* nauplii survival during the whole feeding process. The reservoir can be closed with a lid or left open, and remains at atmospheric pressure at all times. The pump is triggered by a signal coming from the main module and sends the liquid to the module’s base for delivery (Figure 2, D).

### Modules construction

Modules have been designed with Fusion 360 (Autodesk). Custom mechanical parts have been 3D-printed, mostly with PLA (on an Anet A8 printer) and for some parts in PETG (on a Prusa Mk3 printer). The lid of the liquid reservoir’s skirt has been laser-cut (5mm PPMA on a Full Spectrum Laser Hobby machine). Mechanical assembly has been realized with M3 metallic threaded inserts.

The electronics was custom-made and based on inexpensive, well-documented microcontrollers (Arduino Nano v3.1). For the liquid food module, a dedicated Printed Circuit Board (PCB) has been designed (Eagle, Autodesk) and manufactured (PCBWay) to reduce the footprint, avoid mistakes during soldering and ease mounting. The system is able to detect which module is mounted by means of a resistance specific to each module that creates a voltage divider (see Supplementary Materials).

### Calibration

A dedicated setup has been developed for calibrating the solid and liquid food modules, comprising a rigid arm holding the dispenser and a scale (OHAUS PA2102C) as illustrated on Supplementary Figure 1. Both the dispenser and the scale where computer-controlled, and delivered amounts were recorded during series of activation (Supplementary Movies 1 and 2). The same system was used to determine the system’s accuracy with runs of 50 trials of random duration, similar to the human accuracy tests.

### Fish room testing

The system has been tested in the aquatic facility of IBPS (*Sorbonne Université*, approx. 1.100 tanks, 15.000 fish). The experiments were made in agreement with the European Directive 210/63/EU on the protection of animals used for scientific purposes, and the French application decree *Décret 2013-118*. The aquatic facility has been approved by the French *Service for animal protection and health*, with the approval number A-75-05-25.

A simple, preliminary prototype for dispensing solid food with vibration, which was non-modular, without visual feedback and without NFC reader has been routinely used from Sept. 2017 to Sept. 2018. The final version of the system has been used on a daily basis since September 1^st^ 2018.

The survival rates of rotifers (*Rotifera*) and brine shrimps (*Artemia Salina*) have been estimated with visual inspection under a binocular. Movies of rotifers and *Artemia* nauplii have been processed to extract individual trajectories (Figure 3, Supplementary Movie 3 and 4) with a novel cross-species tracking software (*FastTrack*) that will be published elsewhere.

**Figure 3.**
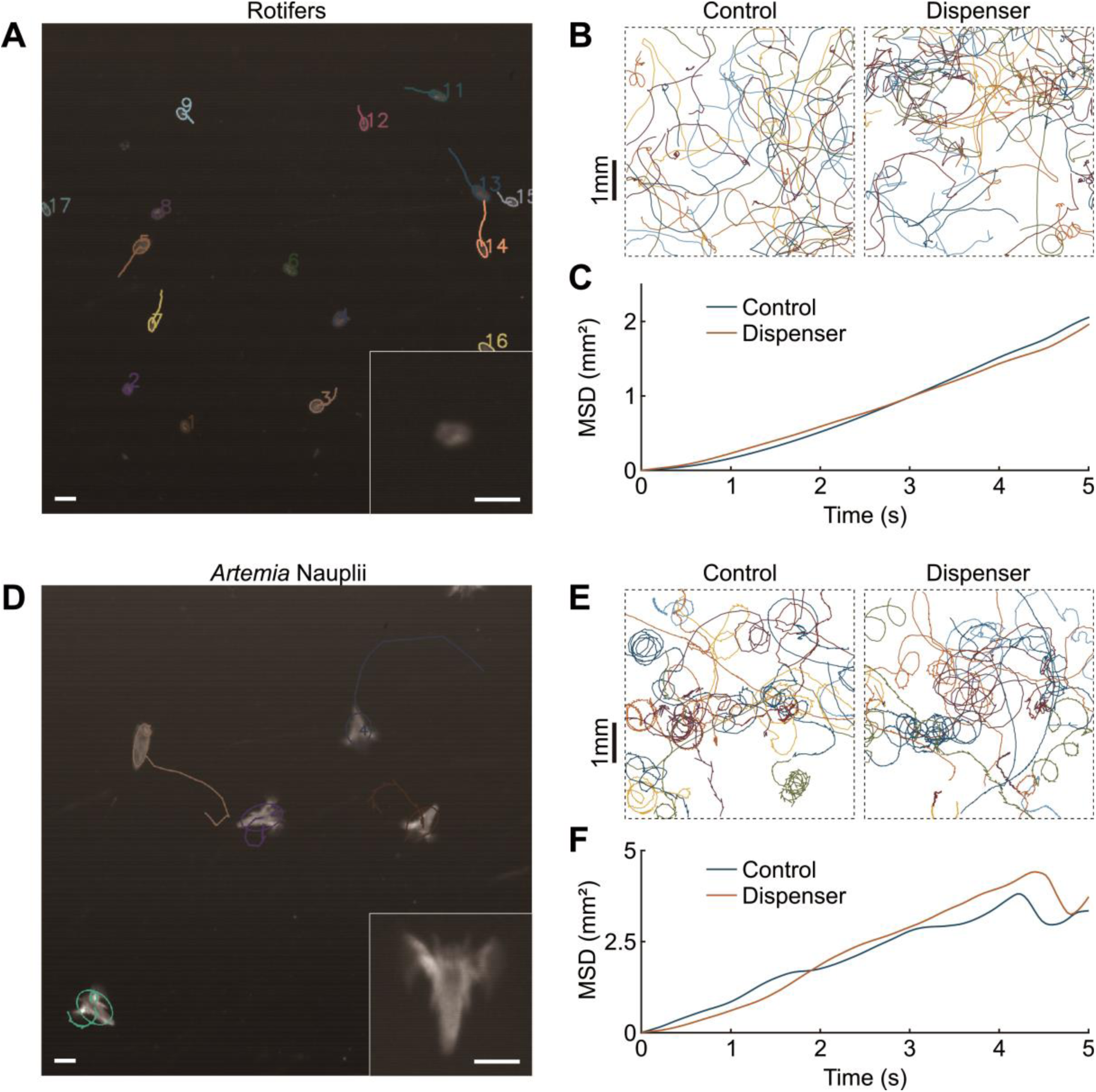
Estimating micro-organisms motility after passing through the dispenser. (**A**) Tracking of a movie of rotifers under a binocular. *Inset*: Blow-up of a single rotifer. Scale bars: 100µm. (**B**) Trajectories and (**C**) *Mean Square Displacement* (MSD) of moving rotifers in the control condition and just after delivery with the dispenser. (**D**) Tracking of a movie of *Artemia* nauplii under a binocular. *Inset*: Blow-up of a single nauplius. Scale bars: 250µm. (**E**) Trajectories and (**F**) MSD of moving *Artemia* nauplii in the control condition and just after delivery with the dispenser.

### Human accuracy tests

Human subjects were composed of two groups: staff of the fish facility who feed zebrafish more than twice a month (trained group, n=6) and people not working in a fish facility selected at random in the population (random group, n=27). Subject from the random group declared not to suffer from a musculoskeletal disorder. All subjects were aged between 18 and 62 and had no information about the setup or the purpose of the experiment beforehand, except that it would last approximatively 30 minutes. Experiments were not remunerated.

The tests were performed in a dedicated room containing only a table with the setup and a chair (Supplementary Figure 5). Subjects were instructed to enter the room alone, close the door, sit down and follow instructions on the screen. All steps of the protocol are detailed in the Supplementary Materials. The subjects had to perform successively four parts with 50 trials each, preceded by short training phases of three trials. Parts consisted in 1) pressing a button for a given duration with immediate visual feedback, 2) pressing a button for a given duration without visual feedback, 3) delivering given amounts of powder with a spoon and 4) delivering given amounts of liquid with a wash bottle.

The program controlling the screen, scale and button has been custom-made and written in C++ (Qt 5.8). The scale (OHAUS PA2102C) was controlled *via* a serial RS-232 connection and was blinded in a black box such that the subject could not see the LCD screen of the scale. The button was a red pushbutton (normally open switch without latch) mounted on a black plastic box containing a microcontroller (Arduino Nano v3.1) and linked to the computer *via* USB. The powder (Sucrose, Merck 84097-250G) was disposed in a beaker with a small spoon. The liquid (water with a blue dye, Indigo Carmine, Merck 57000-100G-F) was disposed in a 500mL wash bottle. A supplementary 250mL bottle was also provided in case the subject had to refill the wash bottle during the course of the experiment.

### Analysis

All data from the system calibration setup, the fish room tests and from accuracy tests have been processed with custom scripts in Matlab (R2018a, The MathWorks).

## Results

Operation of the system in a fish room is presented in Supplementary Movies 1 and 2.

### Live food survival

Live food is generally preferable to purely artificial diets^12,13^, as live feeds possess balanced nutritional profiles^9^, are visually and chemically attractive to fish, and are highly digestible^14^. It also contributes to the animal’s welfare in captivity with the ability to actively hunt and express natural feeding behaviors^15^. It is also mandatory according to the European regulation.

We estimated the survival rates of two standard live feeds, rotifers and *Artemia* nauplii. In control solutions, all animals were moving normally (Figure 3, B and E). After going through the liquid food module, all rotifers were moving in a similar fashion (Figure 3, B) and their observed survival rate was 100%. For *Artemia* it appeared that 5 to 10% of the nauplii died during delivery, probably crushed while passing through the pump. In addition, 20 to 25% of the nauplii were stunned and stopped moving for a few tenth of seconds before resuming normal motion. Trajectories of moving animals were very similar to control (Figure 3, E). We further quantified the motion of rotifers and *Artemia* nauplii before (control) and after delivery by plotting the Mean Square Displacement (MSD) over all trajectories as a function of time (Figure 3, C and F). This is a classical way of characterizing a diffusive process^16^: a straight line indicates diffusive motion, and the slope is equal to 4 times the diffusion coefficient. The coincidence of the MSD curves in both conditions for rotifers and brine shrimps indicate that the dispenser had no measurable effect on the dynamics of the motile micro-organisms, as compared to control.

In practice, Zebrafish ingest inert nauplii as well and no waste is left after a few minutes. Dead and stunned *Artemia* should in principle sink to the bottom of the tank but the tumult created by fish agitation in presence of food scatters the nauplii everywhere in the tank. Careful observation of fish behavior during feeding with the liquid food module make us suggest that such a 3:1 mixture of mobile and inert nauplii could be *in fine* beneficial to the fish, as they are forced to search for food in various places of the tanks and may adopt richer hunt strategies regarding competition with mates.

### Comparison of human and machine accuracy

We developed a psychophysics setup (Supplementary Figure 5) to measure the average accuracy of humans in delivering powder with a spoon and liquid with a wash bottle, and pressing for a given amount of time on a button, with or without visual feedback. These four tasks encompass all reasonable scenarii for manual or manually-assisted delivery. In all tasks the subjects were asked to deliver random integer quantities between 1 and 50, corresponding to a number of animals in arbitrary units (see Supplementary Materials).

Experiments on humans revealed a generally poor accuracy (Figure 4, A-D). Surprisingly, we observed no difference between the performance of the trained and random groups, so we pooled all the data for analyses. Errors, quantified as the delivered amount minus the target amount, were symmetrical and Gaussian-distributed (Figure 4, G) with a dispersion that had little dependence on the target amount. The standard deviation of errors with liquid and solid were very high with 7.8 and 8.2 individuals respectively. These values can be considered too large for research-grade rearing; indeed, undernutrition may lead to developmental issues and fertility losses while overfeeding rapidly degrades water quality. Subjects performed slightly better in measuring time with a button, presumably because the mechanical action is reduced to its minimum, with a standard deviation of 5.0 individuals. Having a visual feedback of the elapsed time further improved accuracy, with a standard deviation of 2.8 individuals. Though this is presumably the best accuracy human subjects can achieve, transposed in a fish room that would require that the operator focuses on a screen while feeding.

**Figure 4.**
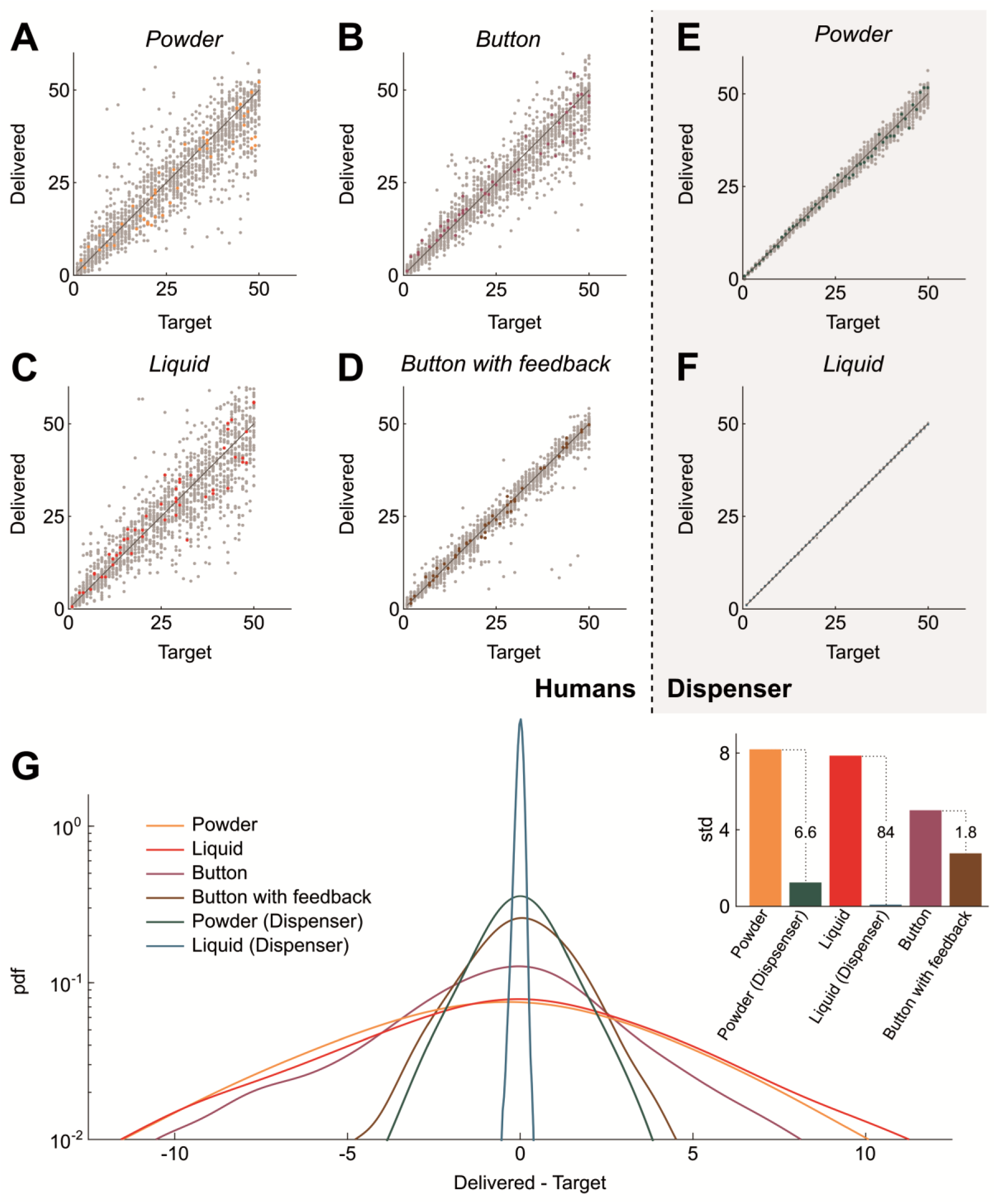
Comparison of Human and dispenser accuracy. (**A**-**D**) Quantity delivered by humans as a function of the target quantity in the four tested conditions: powder (**A**), liquid (**B**), button (**C**) and button with visual feedback (**D**). Data points from all subjects (n=33) are shown in gray and data from one individual chosen at random are highlighted in color. (**E**-**F**) Quantity delivered by the dispenser as a function of the target quantity for powder (**E**) and liquid (**F**) media. Data points from an equal number of runs (n=33) are show in gray and one randomly chosen run is highlighted in color. (**G**) Probability density functions (pdf) of the difference between delivered and target quantities for the different conditions with Humans and dispenser. *Inset*: Standard deviations (std). Additional numbers on dotted lines indicate std ratios.

In our system, NFC detection allows for the dispenser to know the number of animals in the tank and calculate the delivery time accordingly. The operator thus only manages the correct placement of the dispenser over the tank, and the accuracy is solely set by the dispenser’s own reproducibility. The latter is excellent (Figure 4, E-F) and the standard deviation of errors goes down to 1.2 individuals for the solid food module and as low as 0.3 individuals for the liquid food module (Figure 4, G), yielding respectively 7-fold and 84-fold decreases in standard deviation as compared to human performance for similar tasks.

For liquid feed the fluctuations in microorganism concentration are the main remaining source of inaccuracy, while for solid food errors come from minute irregularities in the flow rate. We also expect that for powders the reproducibility is highly sensitive to hygrometry. We also checked that the reservoir tube has to be tightly fixed to the sheath, unless large errors can appear (Supplementary Figure 5).

### Feeding duration

We measured the average time to feed a complete rack *via* manual feeding (wash bottle for liquid, seed sower for solid food) and semi-automatic feeding in NFC mode (Supplementary Table 4). For unexperienced people, the semi-automatic dispenser slightly increased the feeding time (average +7.5%) with solid food and decreased the feeding time (average −21%) for liquid food. For trained staff, we observed a systematic increase of duration with the semi-automatic dispenser (average +56% for both solid and liquid food).

This discrepancy can be explained by the fact that trained staff have naturally developed stereotyped gestures over the years for optimizing the manual feeding time. These stereotyped gestures are not desirable in general, since they generate and aggravate musculoskeletal disorders. For trained staff, the longer time spent in front each rack with the semi-automatic dispenser is partly compensated by the consequent reduction of food-filling episodes: the liquid reservoir has a maximal volume equal to 16 wash bottles and the tubes containing solid food can be carried along and loaded very rapidly.

## Discussion

Our semi-automatic dispenser is a new solution to the numerous issues raised by the pivotal but tedious task of feeding in fish rooms. It is more advanced and convenient than other semi-automatic solutions we are aware of (some being published^17^, but most are not): it is versatile, reliable, accurate, truly portable, easy to clean and do not soot over time. It has also several assets as compared to the commercial fully-automatic solution since it is low-cost, low-footprint and let the operator at the center of the feeding process such that discrepancies are immediately detected and corrected. Without the burden of constantly measuring quantities the operator can use the feeding time to perform routine visual inspection of fish health or check tanks labels. It is also readily accessible to untrained operators, such as students or seasonal staff, who can feed immediately and without any loss of accuracy.

In our opinion, one of the most important aspect of this work is the leap on the ground of musculoskeletal disorders. As soon as the system has been introduced in the fish room, all our staff members observed a relief in forearm pains during and after feeding. A staff member who was previously unable to feed due to repeated wrist tendinitis is now able to feed again without particular pain. These preliminary observations are still to be confirmed over time and with more users, but it is already very promising and we hope that this work will inspire other faculties and companies to invest in research for improving staff health. One possible way for future research is to relieve efforts on shoulder and elbow with an exoskeleton^18,19^.

## Acknowledgements

We would like to thank Sylvie Authier, Édouard Mazoni and Jacques Gaudumet for their help and feedback during testing phases. This work has been financed by ANR JCJC program (ANR-16-CE16-0017). We also thank the Satt Lutech (http://www.sattlutech.com) for endorsing the project, helping in patent writing and financing patent fees and prototype replication.

## Author contributions

All authors have participated in drafting the system specifications. RC designed and built the modules, designed the human-testing setup, supervised these experiments, and analyzed the data. AB, ST and JM have conducted the tests in the fish room. RC and AM wrote the manuscript.

## 1 General information

### 1.1 Sources of the prototype

A patent covering the present semi-automatic feeding system has been filed at the French *Institut National de la Propriété Industrielle* (INPI) and extended for international protection. Source materials including plans, electronic diagrams and Arduino microcode are available upon signature of a material transfer agreement.

### 1.2 Sources of the psychophysics experiment

The source code of the program used for human accuracy experiments is available under the terms of the CeCILL licence for French regulation and more generally the terms of the GNU General Public License.

All files are available at the following URL: http://www.raphael.candelier.fr/Papers/SeAFooD/ReproPsycho.zip

### 1.3 Electronics

All parts of the system, including the electronics, have been designed to be accessible and customizable by non-specialists. All components have a through-hole packaging and only basic soldering equipment is required for assembly. For simplicity, we choose the Arduino Nano v.3.1 as microcontrollers for the main and liquid food modules. A PCB (*Printed Circuit Board*) has been designed for housing the electronic components of the liquid food module, in the form of a cape for the Arduino Nano.

Modules are automatically recognized when connected to the main module. A resistance is located in each module between the ID and GND pins, which forms a voltage divider with the internal resistance of the A0 analog input pin of the Arduino. According to Arduino specifications this internal resistance is between 20kΩ and 50kΩ, so the possible resistance ranges are listed below:

**Supplementary Table 1.**
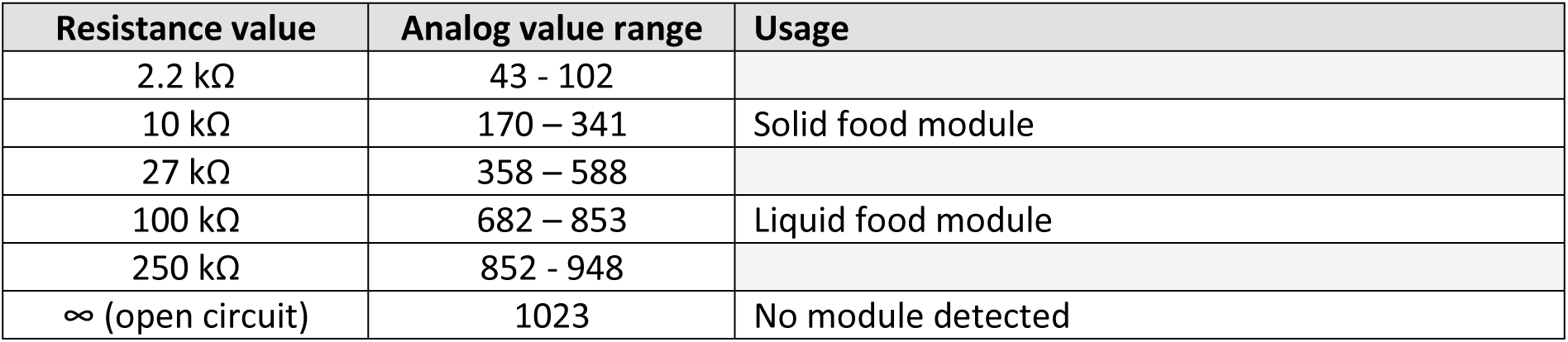
Ranges for the recognition resistance and associated analog values and usages. Grayed cells are reserved for future module development.

### 1.4 Operation and settings

#### 1.4.1 General operation

Both the liquid and solid food module operate at fixed flow rates. The control over delivered quantities is therefore performed by controlling the delivery times *t*_*L*_ and *t*_*S*_, which are related to the number of animals *n* by fixed coefficients *α*_*L*_ and *α*_*S*_:

- For liquid food: *t*_*L*_ = *α*_*L*_. *n*, with by default *α*_*L*_ = 80 ms/fish
- For solid food: *t*_*S*_ = *α*_*S*_. *n* with by default *α*_*S*_ = 35 ms/fish

Note that both *α*_*L*_ and *α*_*S*_ have to be adapted to the food type, the feeding protocol and the fish diet: *α*_*L*_ depends linearly on the concentration in microorganisms (unless the concentration is very high and change the effective viscosity of the liquid medium, but this is not recommended) while *α*_*S*_ depends on the type of solid food and the size of the hole on the delivery tube. The default value of *α*_*S*_ has been optimized for grains 300-500µm in diameter (Gemma Wean 0.3) and a hole of 2.8mm. The coefficients can be changed in the settings menu and are kept in memory even when the device is turned off.

When a solid or liquid food module is mounted on the main module, the device automatically detects which module is mounted and chooses which coefficient *α* to use accordingly.

The system can function in three different modes:

- **Manual mode**. The delivery time is exactly the time during which the trigger is active. The operator thus directly controls the delivered quantity by pulling the trigger for a short or long duration.
- **Fixed mode**. In this mode the amount of delivered quantity is fixed for all tanks. As soon as the trigger is pulled, the device starts to deliver the fixed quantity, regardless of the state of the trigger until delivery is complete. When delivery is over, the system marks a pause of 1 second before being active again. The fixed quantity can be changed in the settings menu. This mode is particularly interesting for tanks with isolated fish (*n*=1) as precise temporal control for one animal is difficult to achieve in manual mode.
- **NFC mode**. In this mode the system has to read first a *Near-Field Communication* (NFC) tag before being active. Once the quantity to deliver is read, it is displayed on the screen and the trigger becomes active. As soon as the trigger is pulled, the device starts to deliver the required quantity, regardless of the state of the trigger until delivery is complete. These steps can be performed in a single movement (see Supplementary Movie 1 and Supplementary Movie 2) since reading the NFC tag is quasi-instantaneous and usually happens before the trigger touches the tank. Once delivery is complete, the system waits to read a different NFC tag before re-activating the trigger.

The operator can easily switch between these three modes with a rotary encoder. The latter is equipped with an RGB LED that shows a different color for each mode (by default: manual=OFF, fixed=cyan, NFC=yellow). The current mode is also displayed on the LCD screen.

Counters of the total number of deliveries (*i.e.* number of tanks) and the total number of fed animals are always displayed on the screen. It is reset to zeros when the device is switched off.

The system is also equipped with a white power LED that illuminates the tanks, allowing the operator *i)* to check that food is actually arriving in the tank and *ii*) to observe the behavior of the animals during feeding. The intensity of this LED can be different at rest and during delivery, and it can be turned off. Many settings, including the coefficients *α*_*L*_ and *α*_*S*_, colors and LED intensity can be accessed and modified *via* a menu interface on the main module. When the button on the rotary encoder is pressed for a one second the systems enters in the *Menu mode* (red color of the rotary encoder by default). Navigation in the menu is achieved with the following commands:

- Clockwise / counterclockwise rotation of the rotary encoder: change selection
- Rotary encoder button: Enter, validation
- Trigger: Cancel, go back or exit the menu

#### 1.4.2 >Menu tree

**Supplementary Table 2:**
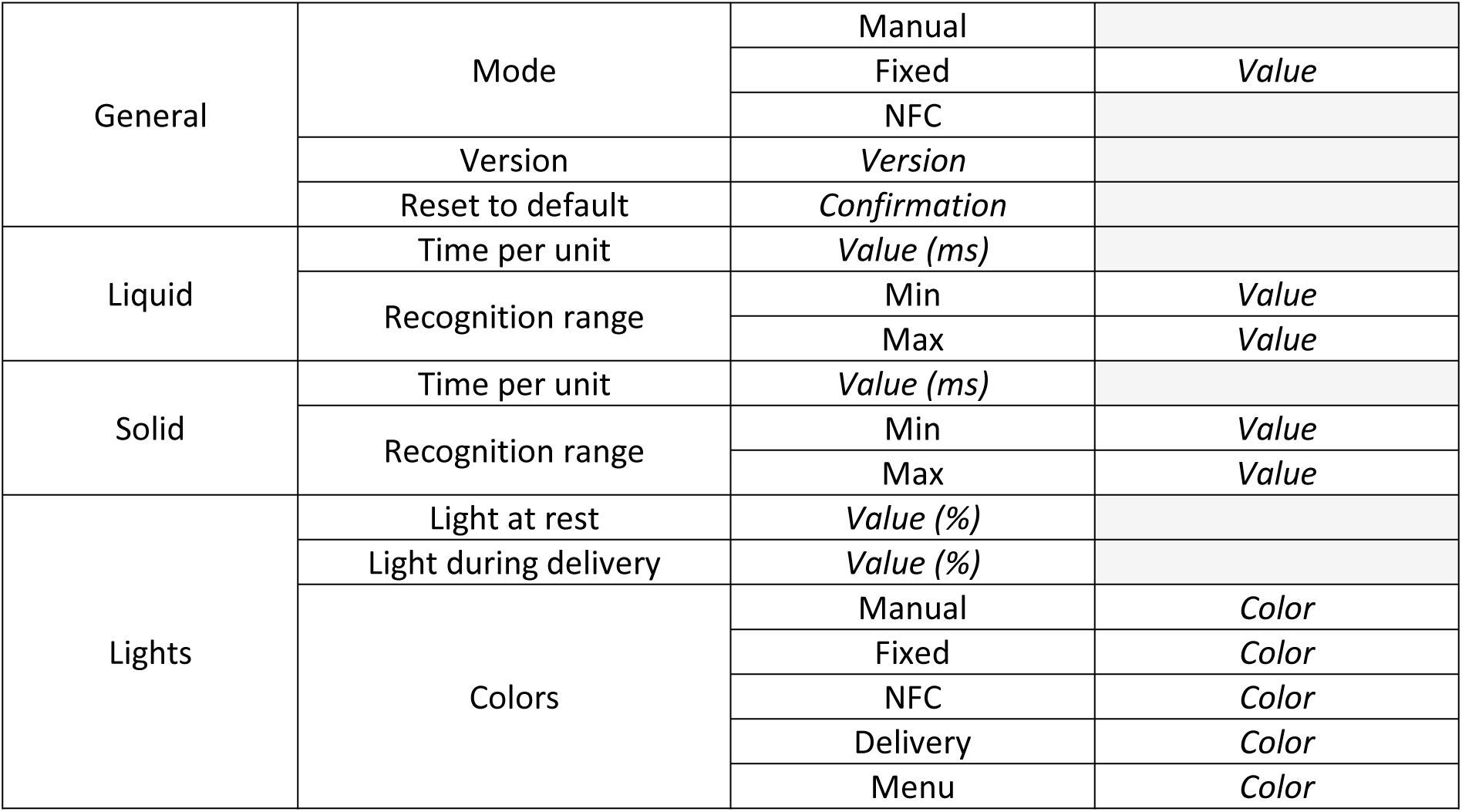
The menu tree, from left to right. Fields in italics are selectable. Color can be either: Red, Green, Blue, Magenta, Yellow, Cyan, White or None.

#### 1.4.3 EEPROM management

The Arduino Nano has an embed EEPROM memory of 1024 bytes. EEPROM is non-volatile, meaning that it can be used to store settings even when the device is powered off. The following table resumes how the 28 first bytes of EEPROM are managed to store settings:

**Supplementary Table 3:** Management of EEPROM memory in the main module.

| Address bytes | Description | Possible values | Default value |
| --- | --- | --- | --- |
| 0 | Version first number | 0 – 255 | 1 |
| 1 | Version second number | 0 – 255 | 2 |
| 2 | Version third number | 0 - 255 | 2 |
| 3 | Version day | 0 - 255 | 17 |
| 4 | Version month | 0 - 255 | 01 |
| 5 | Version year, after 2000 | 0 - 255 | 19 |
| 6 | Mode | 0: Manual, 1: Fixed, 2: NFC | 0 |
| 7 | Fixed value (only for the *Fixed* mode) | 0 - 100 | 5 |
| 10 - 11 | Liquid module detection lower value | 0 - 1024 | 680 |
| 12 - 13 | Liquid module detection higher value | 0 - 1024 | 860 |
| 14 – 15 | Solid module detection lower value | 0 - 1024 | 170 |
| 16 - 17 | Solid module detection higher value | 0 - 1024 | 350 |
| 18 - 19 | Coefficient $\alpha_L$, milliseconds per animal | 0 – 10,000 | 80 |
| 20 – 21 | Coefficient $\alpha_S$, milliseconds per animal | 0 – 10,000 | 35 |
| 22 | LED intensity at rest (%) | 0 – 100 | 1 |
| 23 | LED intensity during delivery (%) | 0 - 100 | 25 |
| 24 | Color at rest, Manual mode | 0: None, 1: White,<br>2: Red, 3: Green,<br>4: Blue, 5: Magenta,<br>6:Cyan, 7: Yellow | 0 |
| 25 | Color at rest, Fixed mode |  | 4 |
| 26 | Color at rest, NFC mode |  | 7 |
| 27 | Color during delivery |  | 1 |
| 28 | Color in the menu |  | 5 |

#### 1.4.4 Nomenclature of the NFC tags: number of animals and cleaning protocol

Data has been written on the NFC tags with the free version of <u>NFCTools</u> and a compatible smartphone (Samsung Galaxy Young 2 SM-G130). The written data is always plain text.

For NFC tags used for feeding, only the number of animals in the tank is written on the tag. So when the main module reads a single number, it is interpreted as a number of animals and the delivery signal is set for the corresponding duration. For instance:

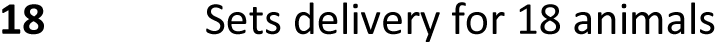

Special diets can be programmed for some tanks only (by punctually changing the value on the NFC tag) or globally (by changing the coefficients *α*_*L*_and *α*_*S*_).

The cleaning procedure of the liquid food module is also defined on a separate NFC tag. When the module reads such a tag, it automatically triggers the cleaning. The protocols are composed of sequences of delivery with 10 second pauses in between. The syntax is as follows:

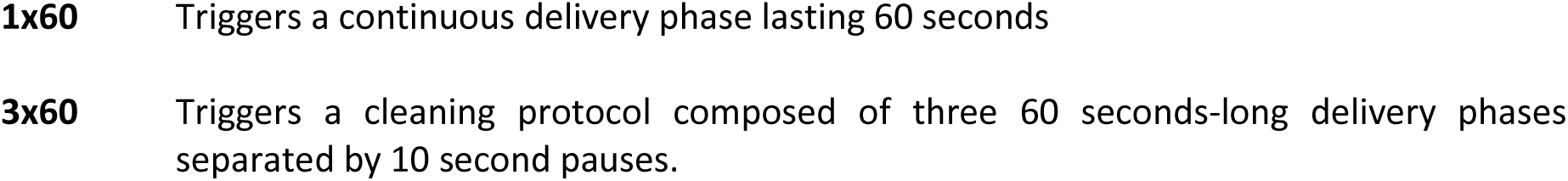

NFC tags can have many shape and materials, and we chose round 25mm-diameter stickers since they are cheaper (small price differences become significant with thousands of tanks). They do not go well in the cleaning machine, so in order to remove and replace them at will we glued it on transparent post-its (see Figure 1.B-C, Supplementary Movie 1 and Supplementary Movie 2) or on thin plastic film usually used for electrostatic stickers. As the information stored on the NFC tags can be re-written endlessly, it is thus easy to reuse it on clean tanks.

## 2 Measuring the system’s accuracy

We used the setup show in Supplementary Figure 1 to measure the system’s accuracy for both the solid and liquid food modules. The scale (OHAUS PA2102C) was connected to the computer *via* an USB to RS232 serial converter. For the solid food tests we used Gemma Wean 0.3 (Planktovie) in a 50mL tube with a 2.8mm drilled hole, and for the liquid food module we used tap water.

**Supplementary Figure 1.**
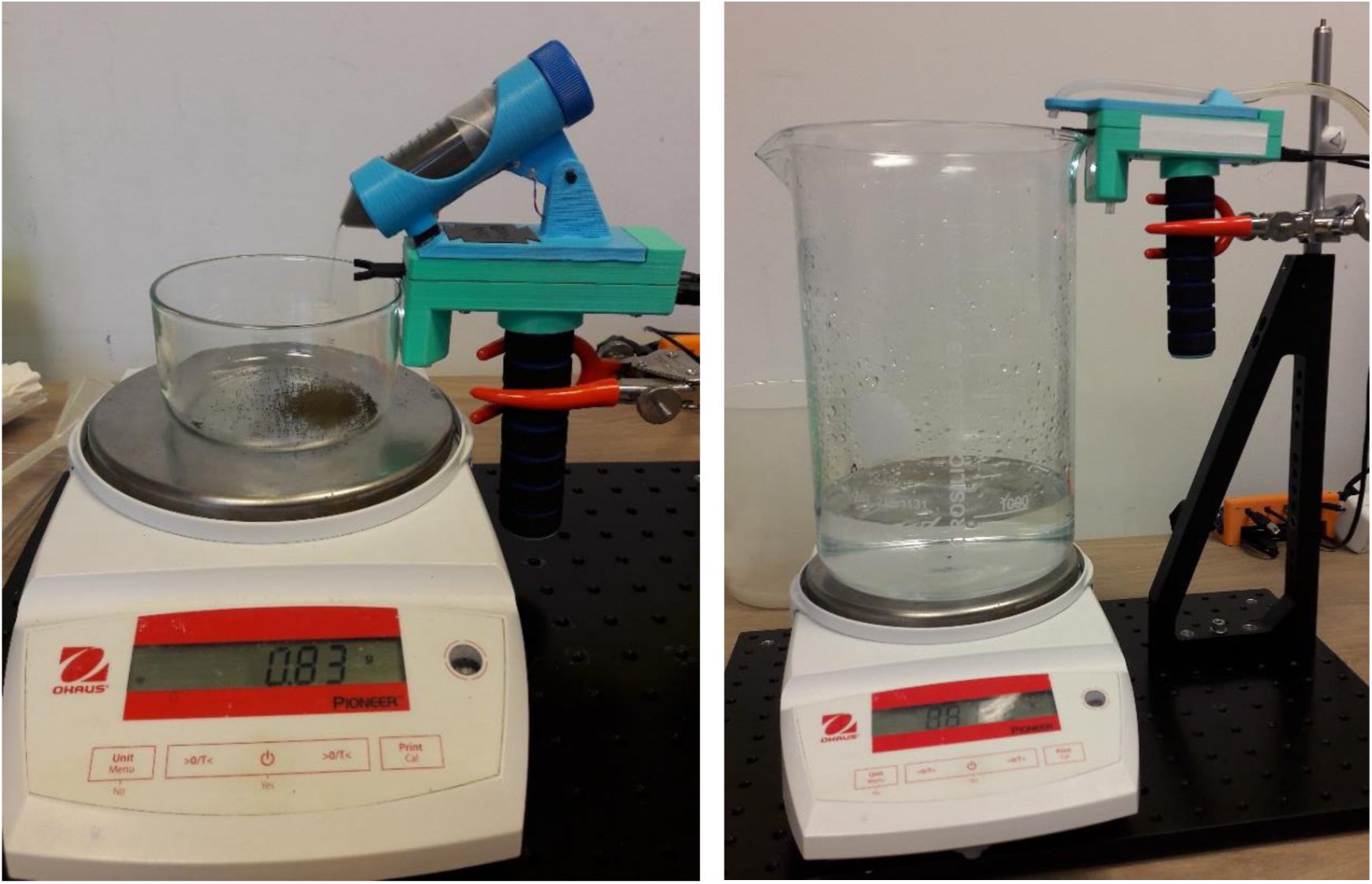
Left: Picture of the setup used to measure quantities delivered by the solid food module. Right: Picture of the setup used to measure quantities delivered by the liquid food module.

### 2.1 Liquid food module

**Supplementary Figure 2.**
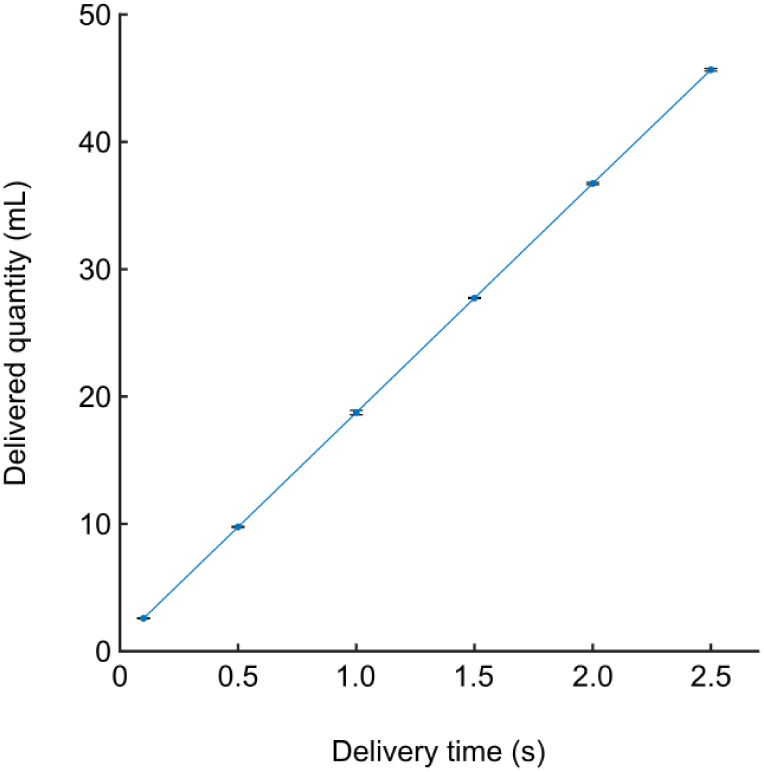
Delivered quantity as a function of the delivering time for the liquid food module. Error bars: standard deviation.

### 2.2 Solid food module

**Supplementary Figure 3.**
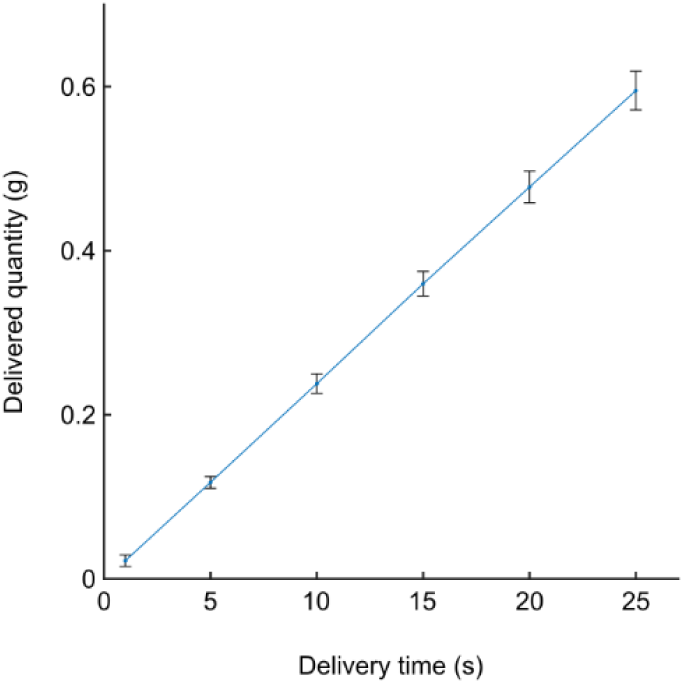
Delivered quantity as a function of the delivering time for the solid food module. Error bars: standard deviation.

In contrast with the liquid food module, several factors can influence the accuracy of the solid food module. For instance, if the containing tube is not tightly attached to the sheath and is able to rotate during vibration the reproducibility is greatly hampered:

**Supplementary Figure 4.**
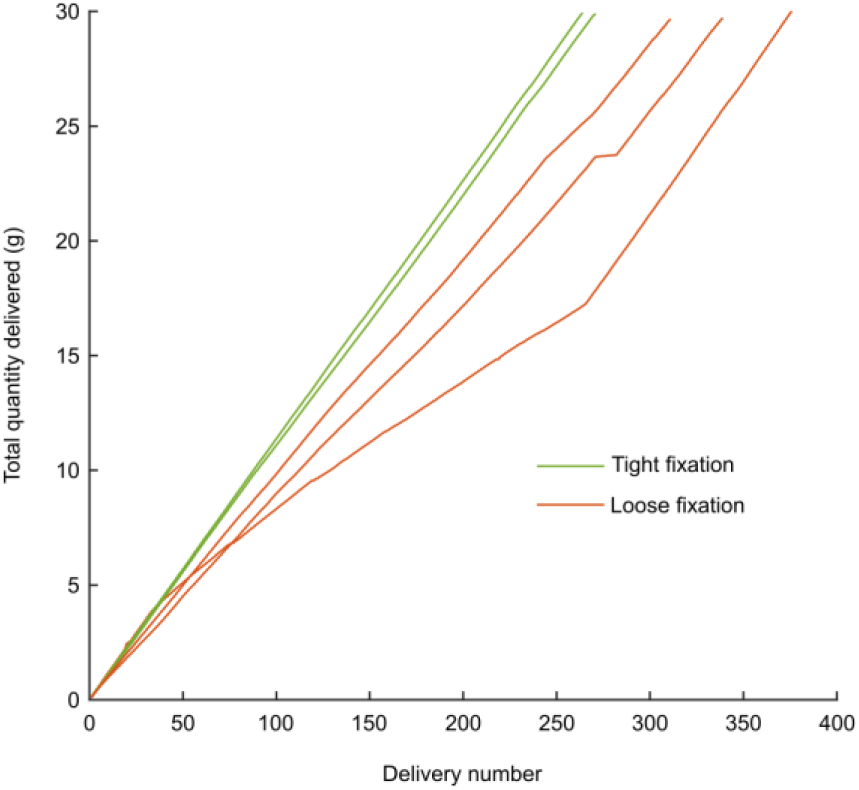
Cumulative quantity of food delivered during runs of 400 trials of 2.0 second of vibration. The containing tube is full (30g) at the beginning of each run and empty at the end. Tight (green) and loose (red) fixation of the containing tube inside the sheath are compared.

Also, controlling the degree of moisture in the powder is essential for accuracy. Water creates bonds between grains that may lead to large aggregates. The latter need a lot of energy to break, so the granular bed is vibration-fluidized with reduced efficiency and severe losses of efficiency can appear. So the food should be kept as dry as possible, which may be difficult in a fish room. We recommend to store the food in a dry place (room with controlled hygrometry or drying box with desiccant).

## 3 Feeding duration

**Supplementary table 4.** Average feeding time (seconds) for rack of 600 fish with an average of 15.4 fish/tank. Fish room staff have fed manually for many years, and used the semi-automatic dispenser for 4 months before the test. Unexperienced operators were volunteer students who had never fed before. Students were shown typical quantities to deliver by trained staff, and they could practice on a few tanks to adjust delivered quantities just before the test. For each condition, 3 subjects were tested on two racks.

|  |  | Fish room staff | Unexperienced people |
| --- | --- | --- | --- |
| Solid food | Manual (seed sower) | 45.6 | 82.7 |
|  | Dispenser (NFC mode) | 71.6 | 89.0 |
| Liquid food | Manual (wash bottle) | 41.2 | 112.5 |
|  | Dispenser (NFC mode) | 64.2 | 88.9 |

## 4 Measuring Human accuracy

### 4.1 Rationale and setup

Our system has a good accuracy over delivered quantities, which we could measure with the automated setup shown in Supplementary Figure 1. But to determine if there is a gain as compared to manual feeding, we had to measure the accuracy of humans in standard feeding tasks. We thus developed a psychophysics setup (Supplementary Figure 5) to measure the average accuracy and reproducibility of human subjects in delivering some liquid with a washer bottle and some powder with a spoon. We also tested human reproducibility on a time-measuring task during which the operator had to press a button for a given amount of time, with or without visual feedback on the elapsed time.

The button was fixed on a small black enclosure containing an Arduino Nano, which was connected to the computer *via* USB. The scale (OHAUS PA2102C) was connected to the computer *via* a USB to RS232 serial converter. The powder was sucrose (Merck 84097-250G) and the liquid was distilled water with a blue dye (Indigo carmine, Merck 57000-100G-F).

Subjects were always asked to deliver an integer quantity between 1 and 50. We used arbitrary units in all parts of the experiment, and the following coefficients (unknown to the subject) were used:

- One unit of time was 100ms for tasks with the button
- One unit of mass was 60mg for the solid delivery task
- One unit of mass was 400mg (400µL) for the liquid delivery task

**Supplementary Figure 5:**
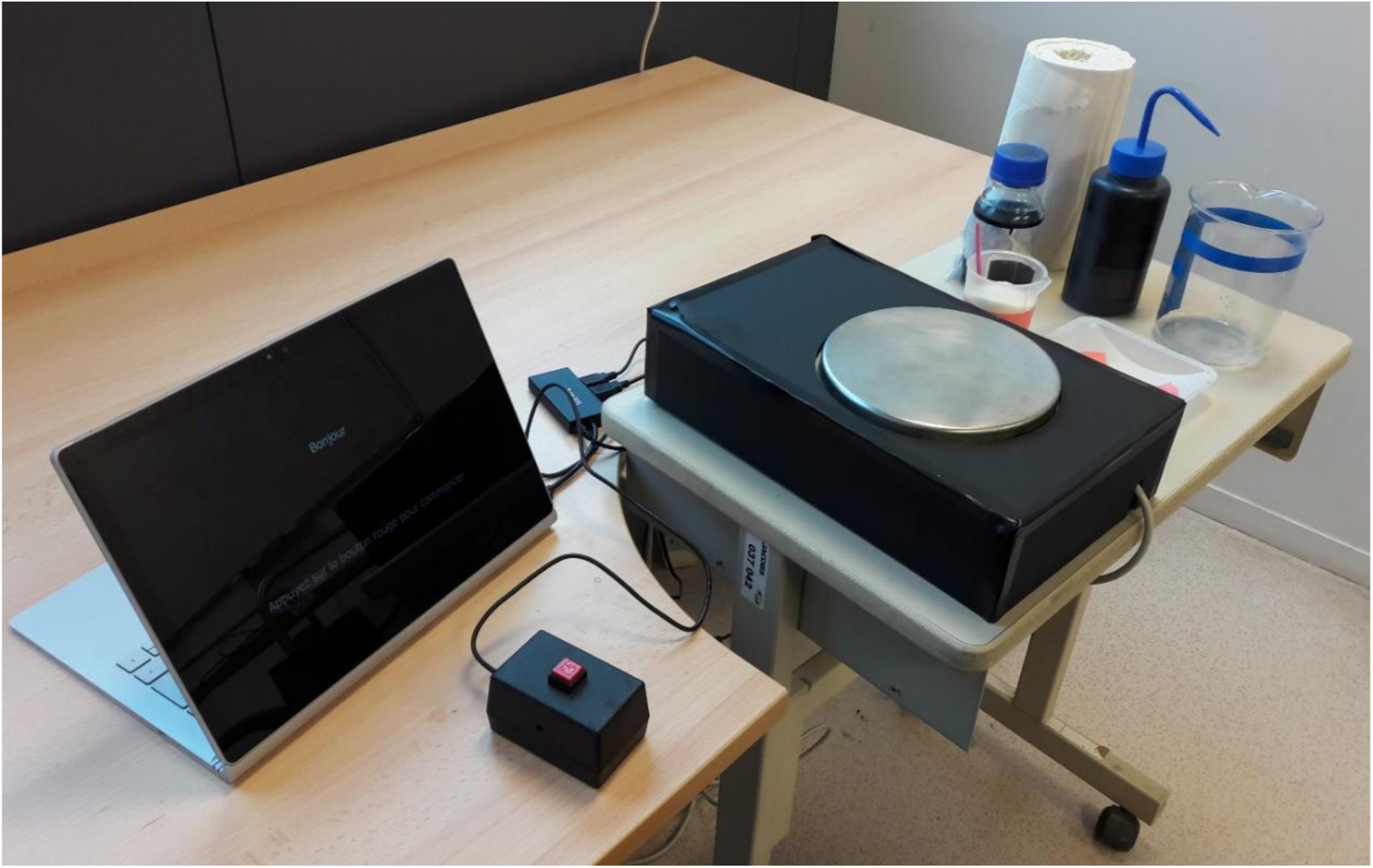
Setup for testing human accuracy in measuring amounts of powder, liquid and time. From left to right: screen, button, scale, powder recipient with spoon (pink), liquid wash bottle.

### 4.2 Step-by-step description

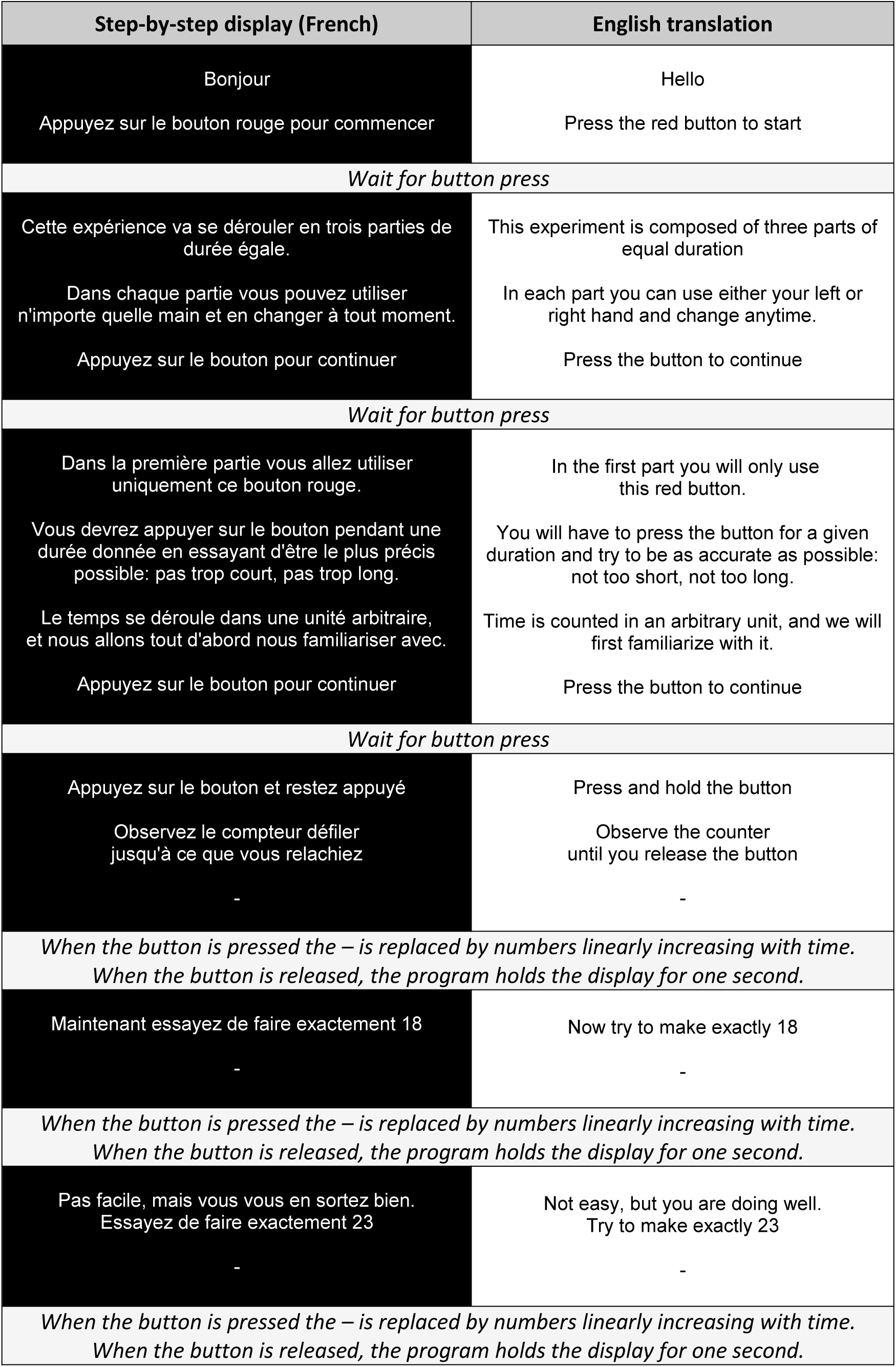
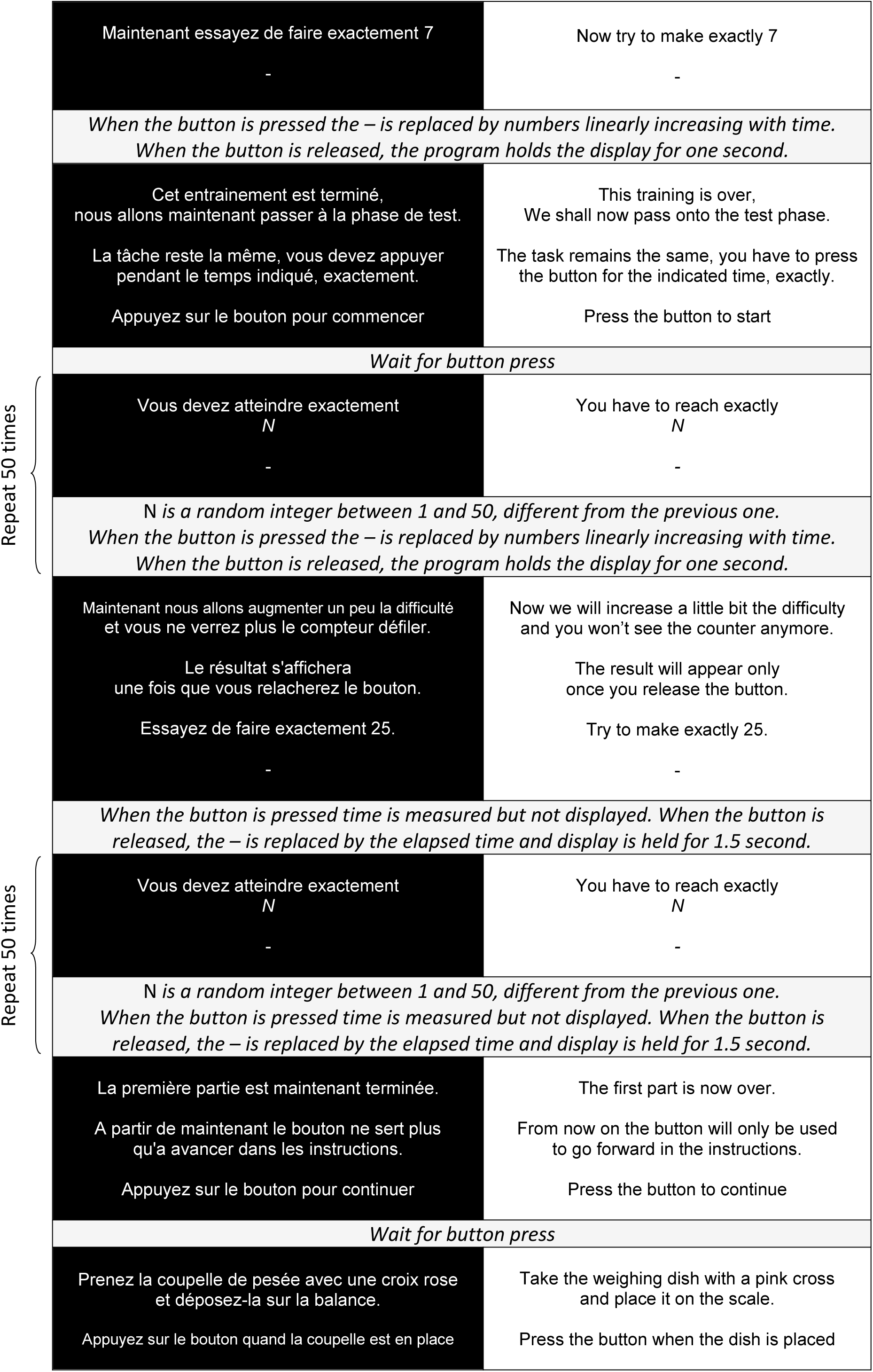
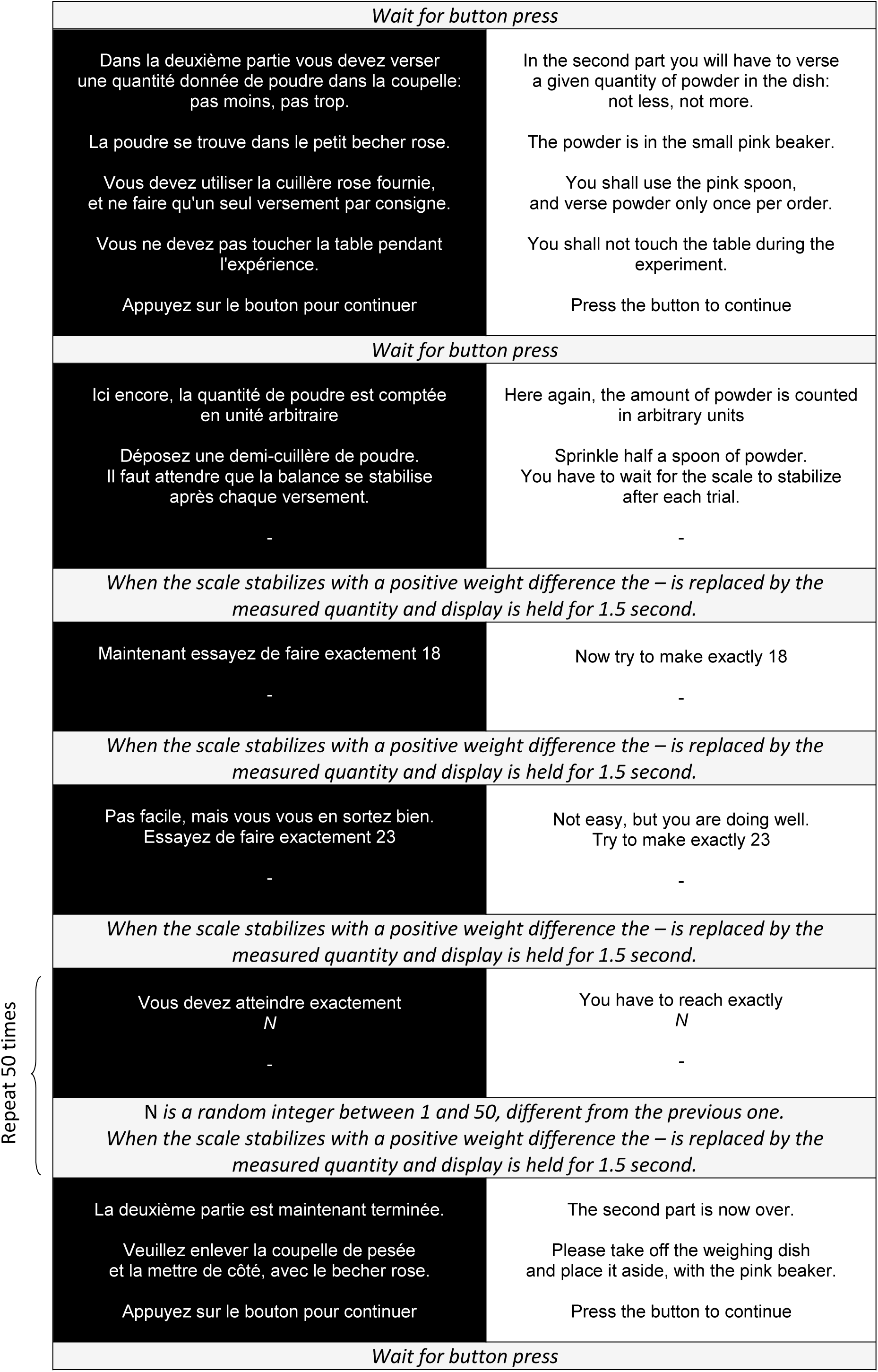
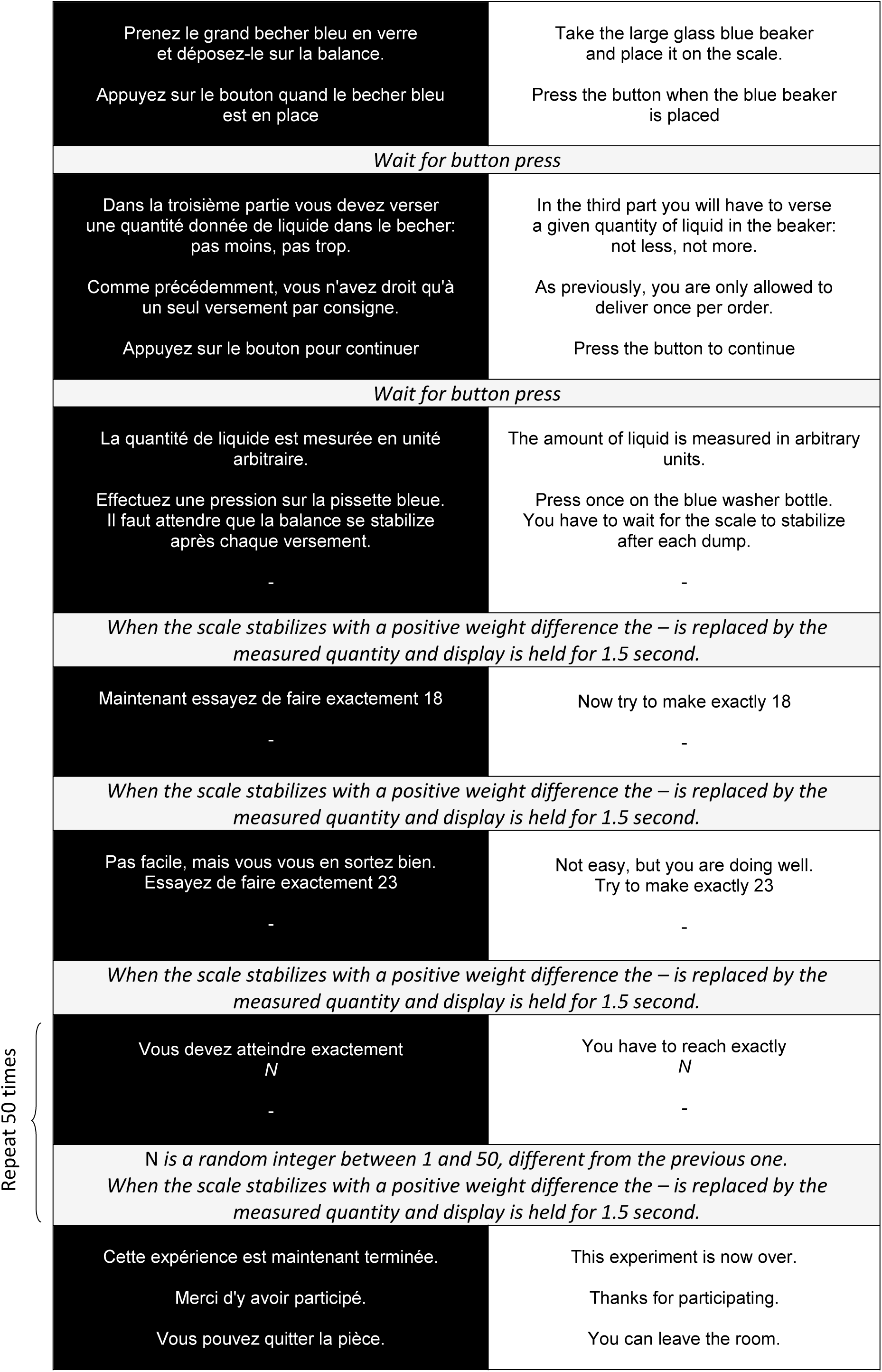

## 5 Supplementary Movies

### 5.1 Supplementary Movie 1

**Solid food module**. 1) Mounting of the solid food module (blue and black) on the main module (cyan). 2) Calibration setup, during a calibration experiment. The system is computer-controlled and delivers for random, uniformly distributed durations while recordings are performed with a scale. 3) Close-up view of the hole upon delivery. 4) Delivery in a tank of a fish room, in NFC mode. 5) Workflow on successive tanks.

### 5.2 Supplementary Movie 2

**Liquid food module**. 1) Mounting of the liquid food module (blue and black) on the main module (cyan). 2) Calibration setup, during a calibration experiment. The system is computer-controlled and delivers for random, uniformly distributed durations while recordings are performed with a scale. 3) Close-up view of the tube end upon delivery. 4) Delivery in tanks of a fish room, in NFC mode. 5) Workflow on successive tanks.

### 5.3 Supplementary Movie 3

**Tracking rotifers.** 1) Motion of rotifers in the control condition. 2) Same movie, with tracking overlaid.

### 5.4 Supplementary Movie 4

**Tracking *Artemia* nauplii.** 1) Motion of brine shrimps in the control condition. 2) Same movie, with tracking overlaid.

